# Genome analysis of two *Lactobacillus plantarum* strains, LLY-606 and pc-26, for evaluating their potential as probiotics

**DOI:** 10.1101/841163

**Authors:** Sudun, Lifeng Liang, Zhenzhen Li, Qingnan Lin, Can Peng, Xuejing Lei, Songling Liu

## Abstract

*Lactobacillus plantarum* was not only one of the most popular probiotics, but also one of the most versatile lactic acid bacteria. *L. plantarum* LLY-606 and *L. plantarum* pc-26 are strains isolated from human gut that are intended to be explored as probiotics. In this study, the genome sequences of LLY-606 and pc-26 were sequenced, and multiple genes related to probiotic properties were analyzed. First, the pathogenicity of these strains was evaluated, and antibiotic resistance genes were surveyed at the whole genome level to determine their primary safety. And then, genes for stress response, plantaricin (pln) biosynthesis, extracellular polysaccharide (EPS) biosynthesis, and bile salt hydrolase (BSH) were analyzed to evaluate their industrial utilization, adhesive capacity, and survival ability in gut, which were properties fundamental for probiotic strains. The physiological features assured by these genes were assayed *in vitro*. The strains were then evaluated *in vivo* for their ability to lower cholesterol, and they were both found to be effective in improving hypercholesterolemia in golden hamsters. In this study, a genetic pre-evaluation was conducted through genome analysis combined with *in vitro* physiological assay, and the probiotic properties of these strains were verified *in vivo*.

## 1. Introduction

*Lactobacillus plantarum* is one of the most versatile lactic acid bacteria [1]. Unlike most other species that are exclusively found in specific habitats [2, 3], *L. plantarum* is found in a variety of different environments such as plants, gastro-intestinal tracts, food materials [4]. Its ability to adapt to various kinds of environments has endowed it with wide industrial utility and potent probiotic effects on its host. Several comparative genome studies have also demonstrated that *L. plantarum* has extraordinary genomic diversity-this extreme genetic versatility is supposed to be the basis of its vast applicability and probiotic properties [4, 5].

However, this extremely versatile property has made the selection of potent strains with specific properties, such as safety, adaptation to host gastro-intestine, industrial utility and specific probiotic functions, more complex [6]. Genomic analysis has facilitated a more comprehensive characterization of the genetic characteristics of some bacterial strains [7, 8]. Some functional genes related to probiotic properties such as bile salt hydrolase (BSH) [9, 10], extracellular polysaccharides (EPS) biosynthesis [11], and stress responses [12] had been identified and illuminated in large number of strains. Moreover, software for analyzing certain properties that are conferred by multiple and unknown genes, such as antimicrobial activity [13], pathogenicity [14, 15] have been explored. The genomic analysis techniques have allowed researchers to comprehensively pre-evaluate strains for use as probiotics more conveniently.

Genotype-phenotype association analysis was required to confirm the genomic analysis. However, these were mainly focused on specific properties, such as adaptation to [16,17] or colonization [18] in intestinal tract. It is not yet successfully used on the comprehensive evaluation of a probiotic strain. In this study we aimed to explore a probiotic evaluating processes that combines genomic analysis with phenotype validation, using *L. plantarum* LLY-606 (LLY-606) and *L. plantarum* pc-26 (pc-26) as model.

These two strains were isolated from human gut in Guangdong province, China. In our previous studies, their excellent capacity to hydrolyze bile salts was defined *in vitro* [19], which made them candidates for probiotic strains with cholesterol-lowering properties [9]. To gain more insights into genetic elements affecting the probiotic properties of LLY-606 and pc-26, their genomes were sequenced and analyzed. A genetical pre-evaluation approach using genome analysis combined with vitro physiological assay then used to verify the probiotic properties of these strains *in vivo*.

## 2. Materials and Methods

### 2.1 Genome sequencing and assembly

LLY-606 and pc-26 were cultured in MRS broth at 37°C. The total genomic DNA were extracted using a Wizard® Genomic DNA Purification Kit A1620 (Promega, Madison, USA) and sequenced by combination of Illumina HiSeq 2000 (Illumina, Inc., San Diego, USA) and PacBio RSII (Pacific Biosciences of California, Inc., USA) platform. PacBio subreads less than one kilobase were removed. Then the Canu software was used for merger, self-correction and *de novo* assembly [20]. Contig was circularized by an inhouse Perl script, which can recognize the start and the end of a contig. Finally, two approaches were employed to correct the assembled genome by using single base. One is Quiver software, which is based on the PacBio long reads [21]. Another is the combination pipeline of SOAPsnp, SOAPindel (http://soap.genomics.org.cn) and GATK (version 3.8.0) [22], which is based on the Illumina paired-end reads. Both of them take the strategy of calling variants and making corrections. The data reported in this study are deposited in the CNGB Nucleotide Sequence Archive (CNSA: http://db.cngb.org/cnsa), as sequencing project **<u>CNP0000375</u>** with accession numbers **<u>CNS0053740</u>** for LLY-606 and accession numbers **<u>CNS0053741</u>** for pc-26.

### 2.2 Genome annotation

The coding sequences (CDS) were predicted from the complete genome sequences of these two strains by GLIMMER 3.02 [23]. The rRNA, ncRNA and tRNA were predicted by RNAmmer (version 1.2) [24], the pipeline integrated with tRNAscan-SE (version 1.3.1) [25] and Rfam database [26], respectively. For functional annotation, the protein sequences translated by CDS were annotated by the pipeline integrated with COG database [27] and RAST website [28]. Biosynthetic gene clusters were predicted by antiSMASH 3.0 [29].

### 2.3 Phylogenetic analysis

The concatenated single-copy proteins in genome were used for the construction of the phylogenetic tree. MUSCLE (version 3.7) [30] with default parameters was employed to perform multiple sequence alignment of single-copy proteins. Then the phylogenetic tree was built by TreeBeST with maximum likelihood method model and bootstrapping value of 1,000. Final tree visualization was processed by iTol [31].

### 2.4 Analysis of safety-related genes

PathogenFinder was employed to determine whether a strain is pathogenic bacteria [14]. PHASTR web server (http://phaster.ca/) was adopted for rapid identification and annotation of prophage sequences within the whole genomes [15]. The virulence and antimicrobial resistance genes were identified by VirulenceFinder (version 1.5) [32] and ResFinder (version 2.1) [13], respectively. The structure of CRISPRs was determined by CRISPRFinder [33]. The genomic islands were identified by IslandViewer [34]. The gene clusters encoding bacteriocins were predicted at BAGEL3 web server [35] and confirmed by RAST.

### 2.5 Adhesion assay

Human colon carcinoma cells HT-29 were cultured in 1640 medium (Gibco, USA) supplemented with 10% (v/v) fetal calf serum (Gibco, USA) and 100□U/ml penicillin and streptomycin (Gibco, USA). HT-29 cells were then seeded in 12-well plates with glass cover slips at 37□ for 4 days. Bacteria cells from stationary phase were collected and washed in PBS. The bacterial cells dissolved in 1□ml 5×10^8^ CFU/mL 1640 Medium were inoculated into each well containing 1ml 2×10^5^ cell/mL HT-29 cells afterwards. The free bacterial cells were washed to remove unbound bacteria, fixed with methanol and stained with Gram staining. The adherent bacteria were microscopically examined at different magnification levels.

The microbial ability to adhere to HT29 cells was also confirmed by quantifying the percentage of adherent bacteria. Briefly, *Lb. plantarum* strains were added (∼100 bacteria/cell) to HT29 confluent monolayers (10^5^ cells/well seeded in 96-well plates 24h prior the experiment) in PBS as assay medium, and then incubated for 1.5h at 37□ in 5% CO2 atmosphere. After incubation, cells were washed with sterile PBS and lysed by addition of 0.25% Trypsin/EDTA solution. The remaining suspensions with adhering bacteria were serially diluted with saline solution, plated into MRS agar plates, and then incubated for 48h at 37□ under. Adhesion to HT29 cells was calculated as percentage of adhered bacteria correlated to the initial number of bacteria added in each well.

### 2.6 Anti-cholesterolemic activity assays

The experiments were approved by the Institutional Animal Care and Use Committee (IACUC) of Institute of Medicinal Plant Development (approval number: SLXD-17062913). Animals were maintained and experiments were performed in accordance with the Guide for the Care and Use of Laboratory Animals (NIH Publication No. 85–23, revised 1996).

Twelve-week-old Syrian golden hamsters were acquired from Institute of Medicinal Plant Development (Beijing, China). Male golden hamsters were randomly divided into four groups, each group had 10 hamsters. All of them were numbered and fed basic diets for 7 days. The golden hamsters were fed in cages under a stable temperature of 21 ± 2°C with humidity of 45 ± 10% and 12h light-dark cycle. The hamsters had free access to water and their particular food. The four groups were appointed to diets according to the following regimen: (1) Control group: standard food diet; (2) HC group: high cholesterol diet; (3) HC-LLY-606 group: high cholesterol diet supplemented with LLY-606; (4) HC-pc-26 group: high cholesterol diet supplemented with pc-26. During the experimental period of 8-weeks, the HC-LLY-606 group and HC-pc-26 group received 10^8^ CFU of LLY-606 and pc-26 intragastrically every day, respectively. The control group and HC group received an equivalent amount of normal saline orally. After feeding for 8 weeks, the hamsters were anesthetized by injecting 3% sodium pentobarbital (2 ml per kg). Four milliliters blood was taken from each individual. Serum total cholesterol (TC) were quantified by commercial enzymatic assays (BioSino Biotechnology and Science Inc).

## 3. Results

### 3.1 Genome features

LLY-606 genome contained a circular chromosome of 3,194,447 bp and a circular plasmid of 64,808 bp, with a GC content of 44.47% (**Table 1, Figure 1**). pc-26 genome contained a circular chromosome of 3,303,726 bp and two circular plasmids of 61,734 bp and 60,139 bp, respectively, with a GC content of 44.46% (**Table 1, Figure 1**). We identified 3,092 genes in the LLY-606 genome with an average length of 882 bp that occupied 83.73% of the genome. In pc-26 genome, 3,146 genes were identified with an average length of 875 bp and occupied 83.42% of the genome (**Table 1**). Within 3,092 genes annotated from LLY-606 genome, 2,339 (75.64%) genes could be assigned to 22 functional categories in Clusters of Orthologous Group (COG) database. While in pc-26 genome, 3,146 genes were annotated and 2,356 (74.88%) genes could be classified into 22 functional class (**Table S1**). Furthermore, 3,035 and 3,064 genes of LLY-606 and pc-26 genome could be assigned to 273 and 271 SEED subsystems by rapid annotation at RAST database. Two thousand six hundred and two genes are common to the two strains and 264, 274 genes are unique to LLY-606 and pc-26, respectively (**Figure 2**).

**Table 1.**
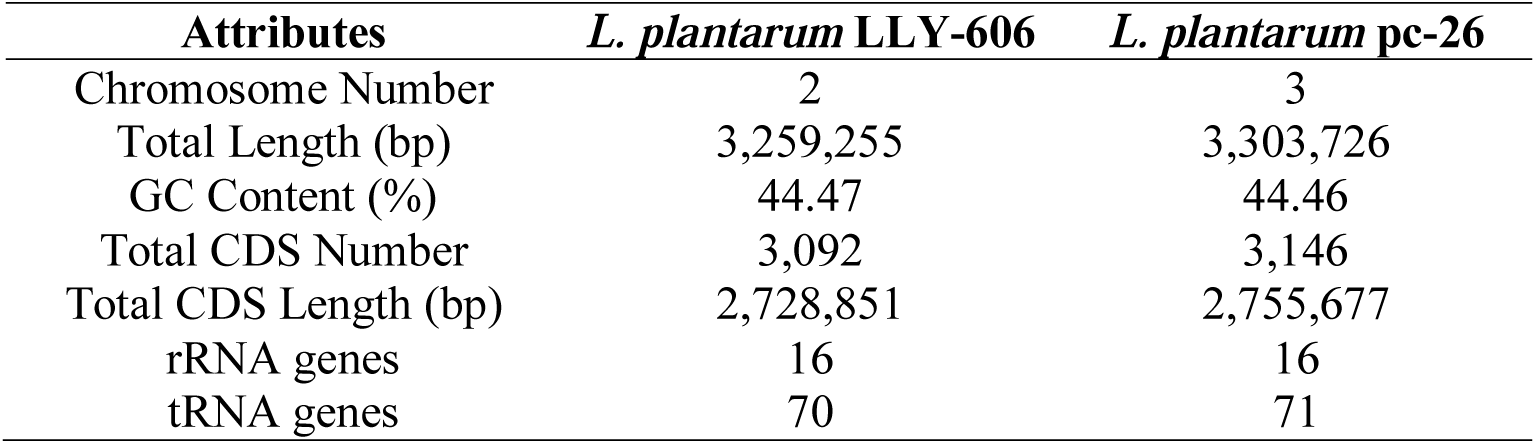
Characteristics of the *L. plantarum* LLY-606 and *L. plantarum* pc-26.

**Figure 1.**
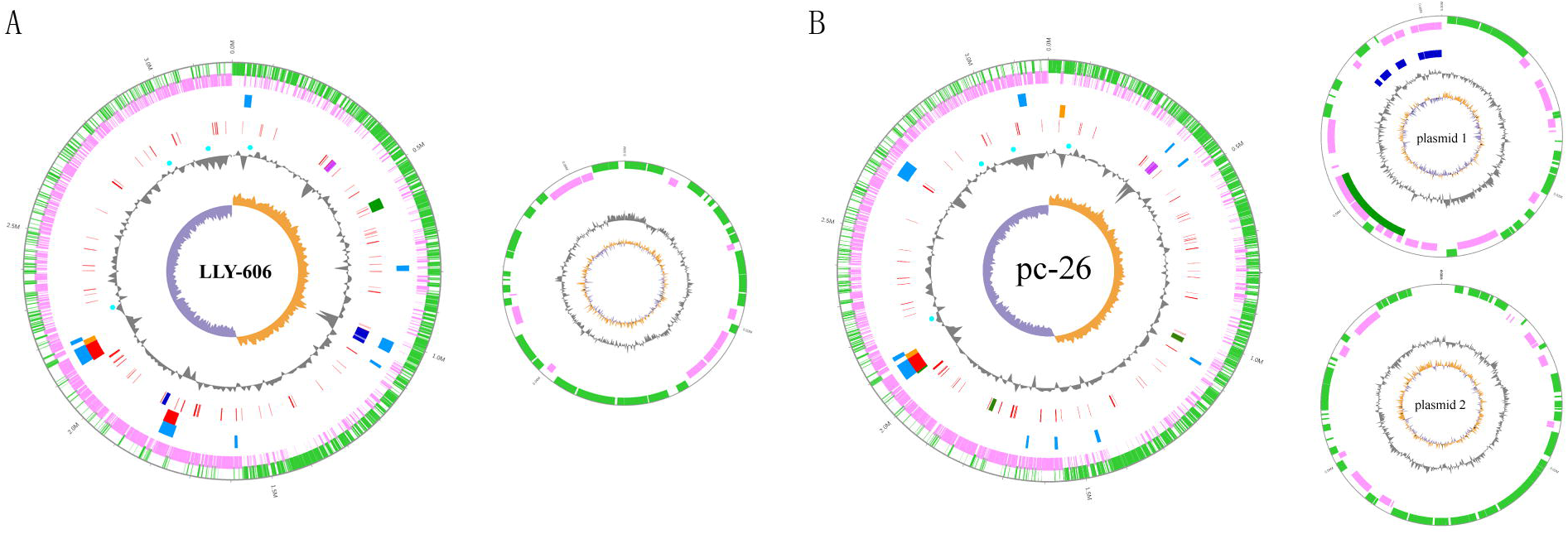
Circular map of the *L. plantarum* LLY-606 genome (**a**) and *L. plantarum* pc-26 genome (**b**). From the innermost, the 1^st^ loop shows the GC skew ((G−C)/(G+C)), where values >0 note in orange, and values <0 note in purple. The 2^nd^ loop shows the (G+C) % deviation from the mean. The 3^rd^ loop denotes *bsh* genes in Cyan bar. In the 4^th^ loop, stress-related genes are in red, capsular polysaccharide biosynthesis genes are in medium blue, plantaricin biosynthesis genes are in magenta. In the 5^th^ loop, intact prophages are in red, incomplete prophages are in orange, questionable prophages are in green. The 6^th^ loop denotes genomic islands in deep sky blue. The 7^th^ and 8^th^ loop indicate the sites of CDS, forward strand are shaded by green and reverse strand are shaded by pink, respectively.

**Figure 2.**
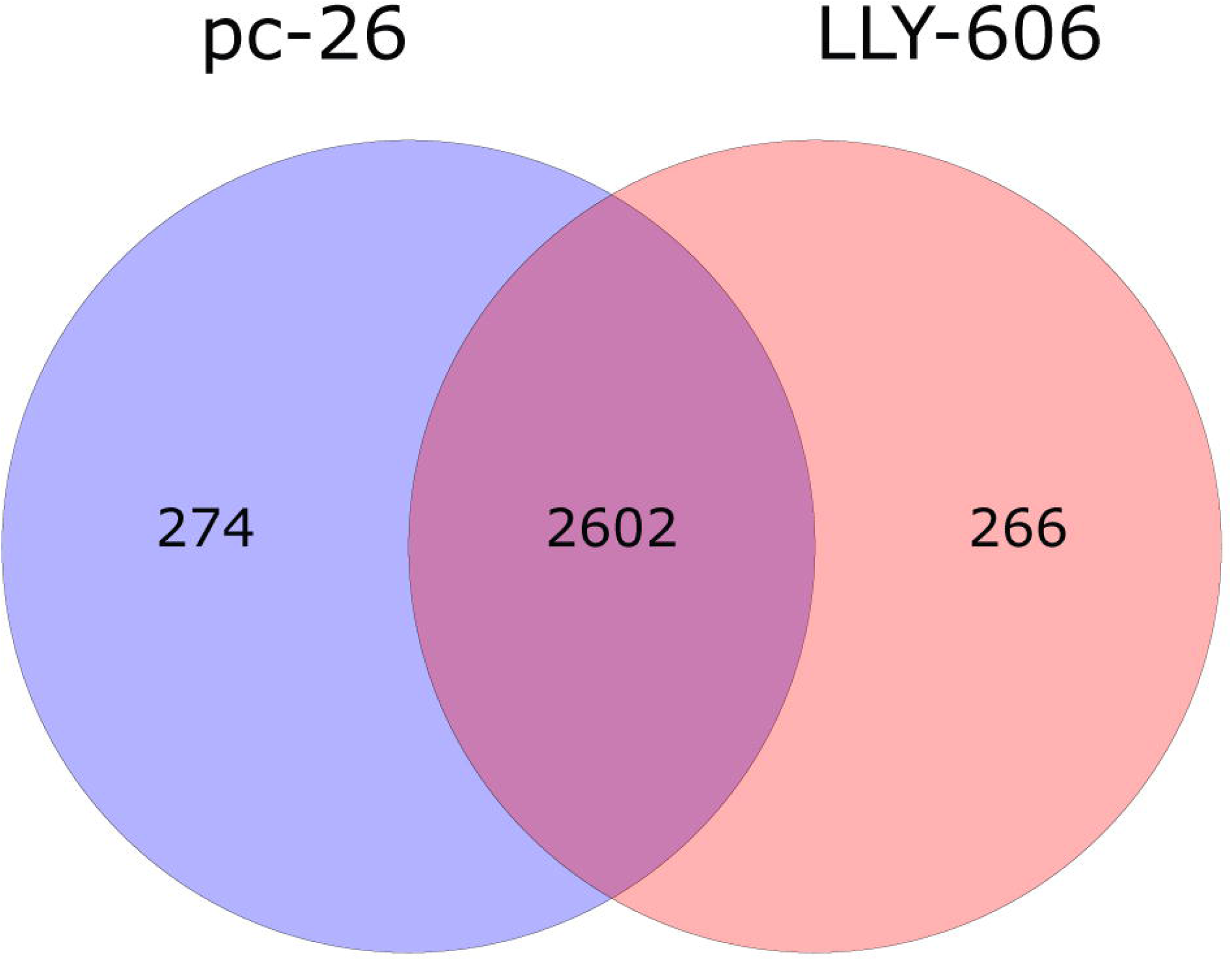
Pan genome of the *L. plantarum* LLY-606 and pc-26.

### 3.2 Phylogeny of the two strains

To reveal the phylogeny of LLY-606 and pc-26, genomes of 56 *L. plantarum* isolate from fermented foods and human intestinal tract were achieved (**Table S2**) and were analyzed. Out of these, 32 strains were isolated from fermented foods, 24 strains are of human origin (feces, gut, saliva), geographically from 19 countries. A phylogenetic tree was constructed by several single-copy ortholog proteins, which no phylogenetic cluster correlate with geographic location or isolation source to be present, even if a small proportion of strains from the similar niches or the same continents were clustered, such as the cluster of Nizo2257, TIFN101, Nizo2258, and the cluster of NCU116, LY-78 and Nizo2801 (**Figure 3**(**a**)).

**Figure 3.**
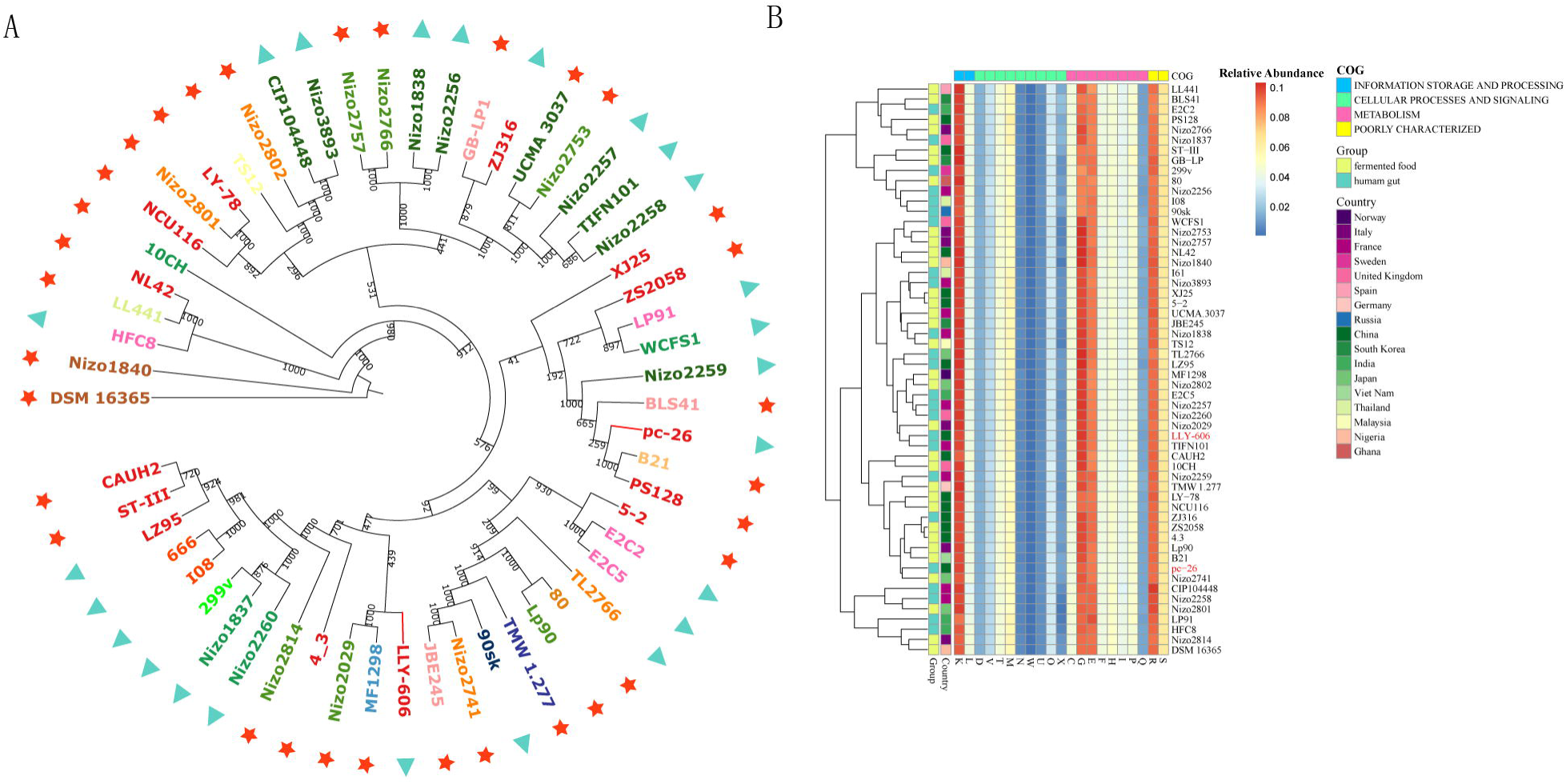
Comparative genome analysis of *L. plantarum*. (**a**) Phylogenetic tree of LLY-606, pc-26 and other 54 *L. plantarum* strains. Strains from fermented foods are represented by red star and strains from human intestinal tract are marked with cyan triangle. Each strain from country of isolation is shaded by its corresponding color in **Figure 3**(**b**). Bootstrap values are shown on each node. (**b**) Heatmap of hierarchical clustering of the COGs categories in the 56 *L. plantarum* strains.

To further investigate their potential habitat-adaptation signatures, the proteins of 56 strains were assigned to Clusters of Orthologous Groups (COGs). As a result, the COGs annotation ratio of all strains was more than 75% (**Figure 3**(**b**), **Table S1**). The enriched categories were related to genetic information storage, processing and biological metabolism, including translation[J], transcription[K], carbohydrate transport and metabolism[G], amino acid transport and metabolism[E]. However, no obvious clustering pattern was identified from living habitats or geographical origin (**Figure 3**(**a**)), and t-test showed that there was no significant difference in all categories from two habitats. Thus, all of these results indicated that the Clusters of Orthologous Groups (COGs) of the *L. plantarum* strains were not correlated with their habitats and geographical origin.

In summary, the strains of *L. plantarum* from two living habitats, fermented foods and human intestinal tract, share similar genome, even if the numbers of each COGs categories are different.

### 3.3 Genome features related to probiotic properties

#### 3.3.1 Analysis of safety-related genes

The genomes of LLY-606 and pc-26 were surveyed using PathogenFinder, and they were genetically negative, indicating that the two strains were not pathogenicity. Antibiotic resistance genes of the two strains were also identified by ResFinder software. And just as the biochemical antibiotic resistance assays [16] showed, LLY-606 and pc-26 were sensitive to the tested antibiotics. Furthermore, virulence factors and CRISPRs were absent in LLY-606 and pc-26 genomes. At last, it was found that LLY-606 contains four prophage elements, including two intact, one incomplete and one questionable (**Table S3**). One region of an intact prophage resembles Lactob_Sha1, which is similar to the prophage region 1 of 5-2 [36]. Another intact prophage resembles Oenoco_phiS13 (**Figure 4**). The strain pc-26 contains five prophage elements, four on the chromosome and one on the plasmid 1(**Table S3**). There is one intact prophage in pc-26, which resembles Oenoco_phiS13 (**Figure 4**). Prophage Lactob_phig1e was absent in LLY-606 and pc-26, which exist in 5-2[36] and WCFS1[37].

**Figure 4.**
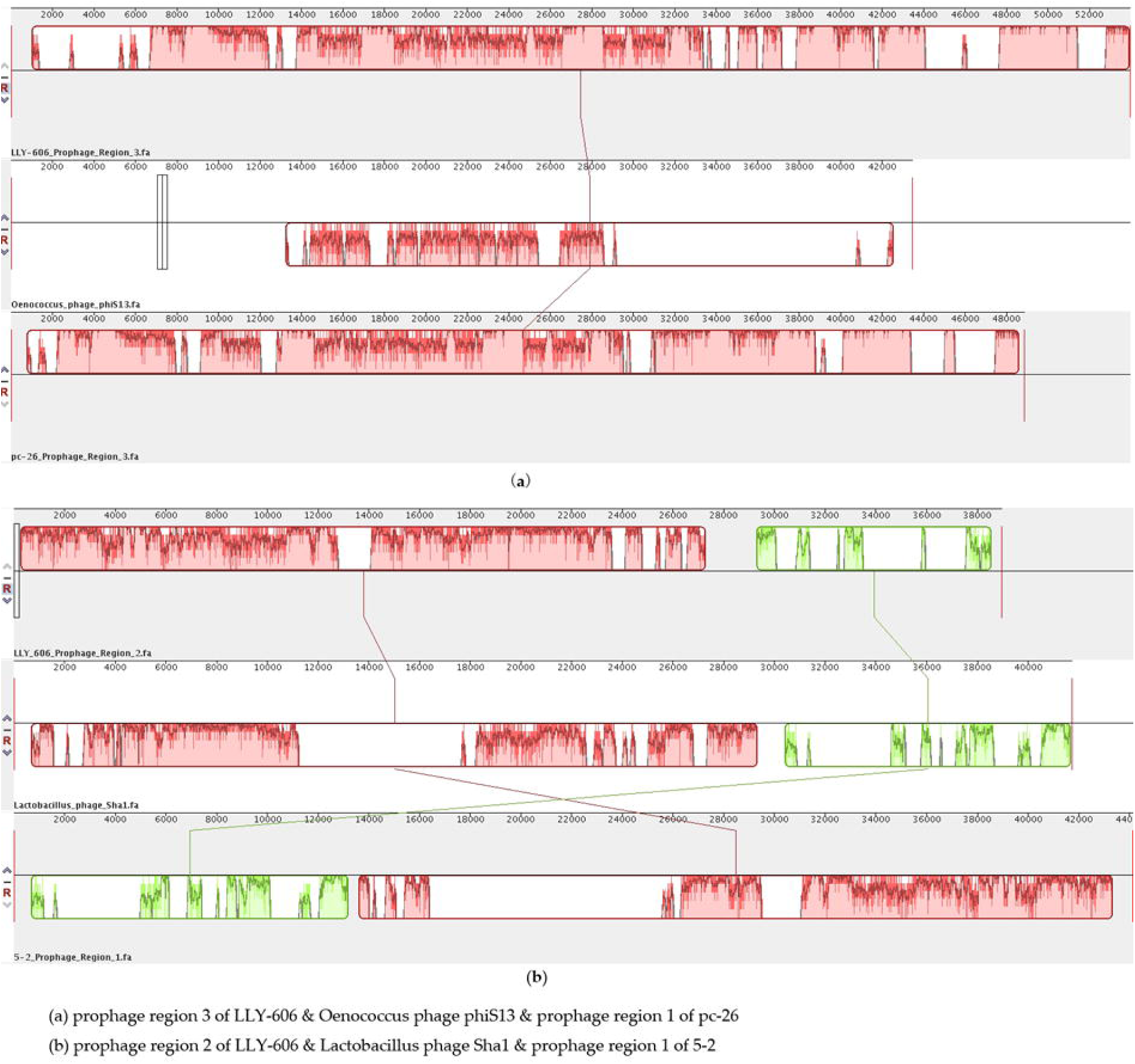
The multiple sequences alignments of prophages of *L. plantarum* and phage.

#### 3.3.2 Stress response

A number of stress-related proteins were identified in both LLY-606 and pc-26 genomes (**Table S4**), including several *Clp ATPases*, universal stress protein (UspA), and *luxS. In vitro* assays were taken out and demonstrated that LLY-606 and pc-26 had a 77% survival rate after exposed to artificial gastric juice of pH 3.0 for two hours [6].

#### 3.3.3 Adhesion ability

In WCFS1 genome, gene clusters of *cps*2 and *cps*4 are reported to encode all functional proteins demanded for capsular polysaccharide synthesis [38]. In LLY-606 and pc-26, we also identified 57 and 35 encoding genes associated with extracellular polysaccharide biosynthesis respectively (**Figure 5**). The *cps*4 in both LLY-606 and pc-26 was similar to *cps*4 in WCFS1. But gene clusters of *cps*2 and *cps*3 in both LLY-606 and pc-26 lost several genes found in WCFS1. The *cps*1 in LLY-606 consisted several genes similar to WCFS1 while *cps*1 was absent in pc-26. Corresponding to the genetical property, an adhesion assay *in vitro* showed that LLY-606 and pc-26 adhered to the surface of HT-29 cells pretty well. The adhesion efficiency of LLY-606 & pc-26 to HT-29 was shown in Figure S1 and Figure S2, respectively. Together, these results indicated that LLY-606 and pc-26 have adhesion ability.

**Figure 5.**
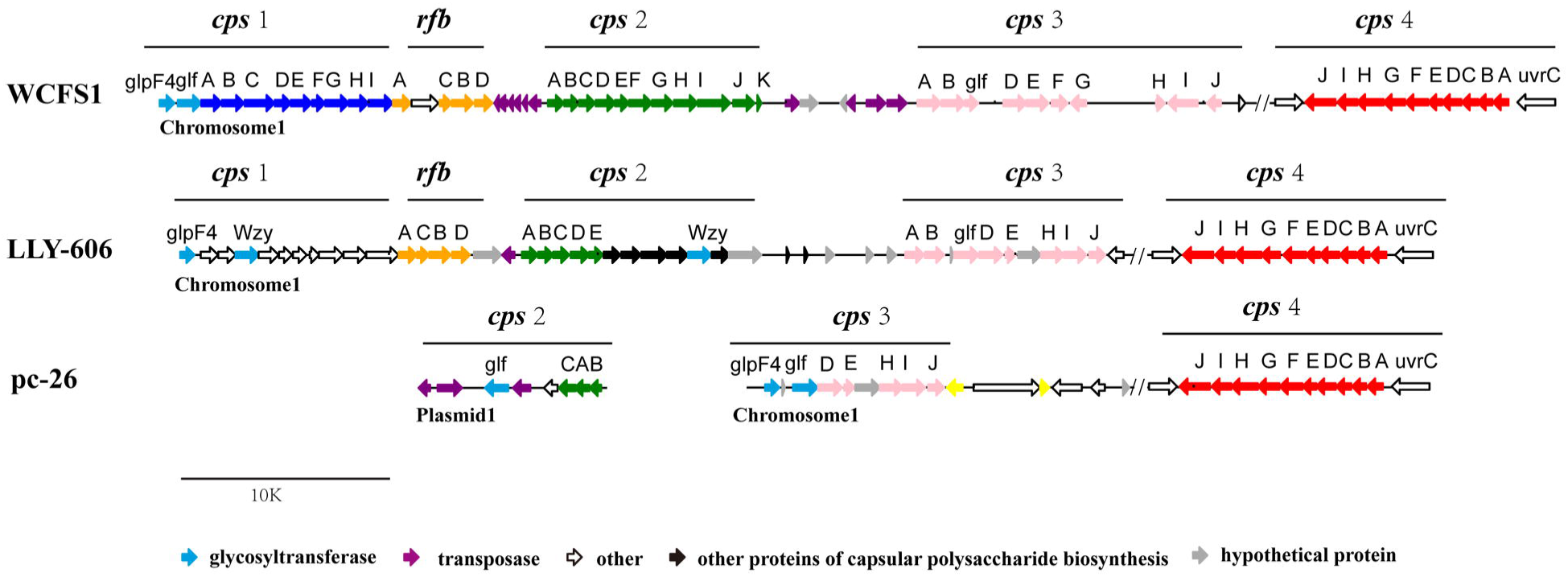
Clusters of capsular polysaccharide biosynthesis genes in LLY-606, pc-26 and WCFS1. Blue, green, pink, red represents gene of *cps* 1-4. The cluster of rhamnose biosynthesis are marked with orange. Purple arrows indicate the transposase, integrase or recombinase. Yellow arrows represent transcription regulator. Black arrows denote genes associated with capsular polysaccharide biosynthesis, which are absent in WCFS1.

#### 3.3.4 Antibacterial activity

It had been demonstrated that LLY-606 and pc-26 both exhibited significant inhibition to *Staphylococcus aureus, Pseudomonas aeruginosa* and *Enterobacter cloacae* [16]. In this study, we identified a pln biosynthesis gene cluster in both LLY-606 and pc-26, which had a structure similar to the cluster found in *L. plantarum* WCFS1 [39] (**Figure 6**). The *pln* gene cluster of pc-26 had 22 encoding genes, including four class IIb bacteriocins (PlnJ, K, E, F), one class IIc bacteriocin (PlnA) and one class I bacteriocin (PlnW) [40]. Complete two-component system (PlnC, PlnD), a response regulator of bacteriocin biosynthesis, was identified in the *pln* gene cluster. The class II bacteriocins are heat-stable and without lanthionine residues [41], which might explain the fact that cell-free supernatant of the two strains still had antibacterial activity after water bath at 85°C for 20min [16].

**Figure 6.**
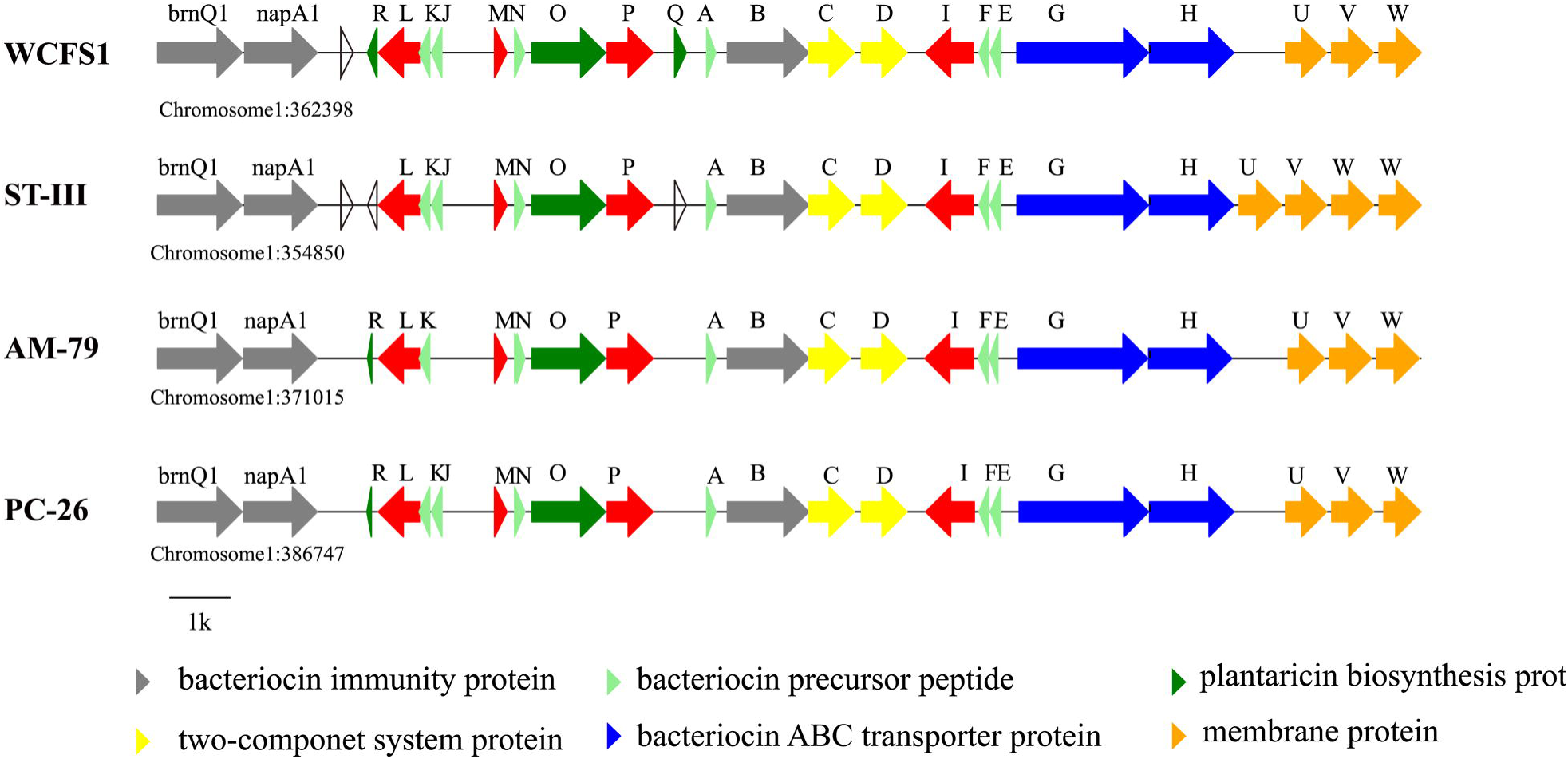
Structures of the plantaricin (*pln*) gene clusters from different *L. plantarum* strains. Each bacteriocin class has different color, the arrows represent *pln* genes.

#### 3.3.5 Bile salt hydrolase

Based on the complete genome sequences, four *bsh* genes were identified in LLY-606 and pc-26, respectively. In *L. plantarum*, some strains have four *bsh* genes, such as WCFS1 [42], ST-III [43], and some have only two *bsh* genes, such as ZJ316 [44]. To further summarize the *bsh* genes of *L. plantarum*, we surveyed 38 complete genomes of *L. plantarum* from NCBI (https://www.ncbi.nlm.nih.gov/genome/genomes/1108). We found that 30 strains had four *bsh* genes, while the remaining 8 strains had less than four, 3 genes for 7 strains and 1 gene for 1 strain. And the locations of these four genes in the genome were generally fixed (**Figure S3, Table S5**).

### 3.4 The anti-cholesterolemic effect of LLY-606 and pc-26

In order to evaluate the anti-cholesterolemic effect of the two *L. plantarum* strains, hypercholesterolemia model was established via high-cholesterol diet (HC) in golden hamsters. The total cholesterol (TC) levels in the serum was tested after 8-weeks administration of the two strains (**Figure 7**). Four groups of hamsters were fed on standard food (Control), high cholesterol diet (HC), high cholesterol diet supplemented with LLY-606 (HC-LLY-606) and pc-26 (HC-pc-26), respectively. Hamsters in the HC group presented elevated TC levels (23.28 mmol/L) compared with the Control group (4.03 mmol/L, p=0.0013). Serum TC levels of HC-LLY-606 group and HC-pc-26 group were significantly decreased (12.31 mmol/L, p=0.0013 and 16.35 mmol/L, p=0.03 in HC-LLY-606 group and HC-pc-26 group, respectively) compared with HC group. Hamsters in the HC-LLY-606 group had significantly lower TC levels (*p*=0.045), compared with the HC-pc-26 group. It was clear that both LLY-606 and pc-26 have a remarkable effect on reducing total cholesterol in serum, and LLY-606 performed better.

**Figure 7.**
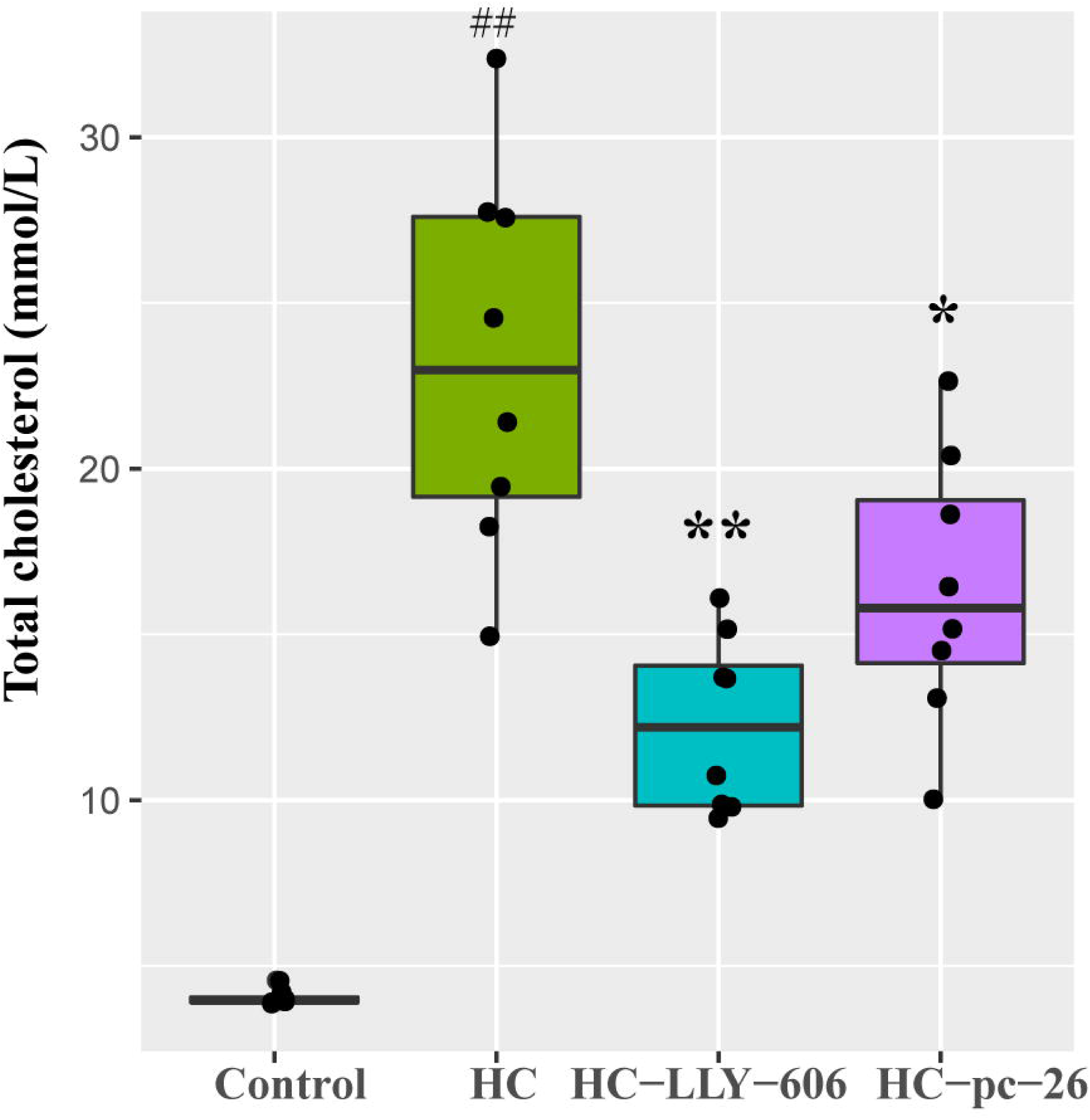
Total cholesterol (TC) levels in the serum of hamsters. Control was the group fed on the standard food diet and HC was of high-cholesterol diet. HC-LLY-606 and HC-pc-26 were fed on the high-cholesterol diet with additive LLY-606 and pc-26, respectively. Values of p were determined using wilcoxon test analysis. ** p□<□0.01 and * p□<□0.05 are versus the HC group; and ## p□<□0.01 is versus the control group.

## 4. Discussion

Although lactic acid bacteria are generally regarded as safe (GRAS), some hazardous conditions were reported [45, 46]. For this reason, safety is still the fundamental requirement for a probiotic [47]. And genome analysis that was based on bioinformatic database and useful software was proved to be the most efficient means to carry out this complex safety pre-evaluation. In this study, pathogenFinder was used to conduct pathogenicity evaluation at genome level. The genetically negative results gave us more confidence to exploit these two potential stains for their probiotic properties. Beside pathogenicity, antibiotic genes, especially those transferable genes were threat to the whole gut microbiota [48]. So antibiotic resistance genes of the two strains were identified by ResFinder software. And just as the biochemical antibiotic resistance assays showed, LLY-606 and pc-26 were sensitive to the tested antibiotics. As antibiotic resistance was wide and dubious properties, biochemical assays directed by pre-evaluation at genomic level was no doubt the first choice for this task. At last, prophage elements were involved in acquisition or loss some genes, which were indications of the stability and origin of the strains. These results provided pieces of evidences for the safety of these two strains, which is a fundamental prerequisite for a potential probiotic candidate [47].

After safety evaluation, it was must be taken into consideration that during both the digestion process in the gastrointestinal tract and during producing, storage, and delivery in industrial processes, probiotics were threatened by environmental stresses, including unsuitable temperature, osmolarity, oxidation and highly acidic conditions [12]. Survival through these harmful conditions and arrival in intestine were essential for probiotic to exert their beneficial effects on their host. Some probiotics are equipped with an array of stress response systems to fight against environmental stresses [49]. Just as expected, a number of stress-related proteins were identified in both LLY-606 and pc-26 genomes which had been demonstrated to play critical roles in acid stress response of *L. acidophilus* NCFM and *L. rhamnosus* GG [50]. However, whether their high survival rates reported previously [16] arose from the stress-response genes and whether the stress-response genes would endow the strain survive these stress conditions, was still needed to be proved in further study.

In addition to high ability to survival hazardous intestinal environments, persistence in intestine mucosa, was also reported to be an essential probiotic property. EPS were reported to play a relevant role in the persistence of the producing bacteria in the intestinal tract [51]. And some EPS locus has been illuminated genetically. What’s more these locus were one of the most variable locus in many genomes [4]. The EPS locus identified in LLY-606 and pc-26 genes, and the adhesive capacity demonstrated in this study indicated their potent to persist in intestinal mucosa to exert their probiotic functions.

As part of their probiotic contribution to the host, the antimicrobial activity by means of bacteriocin production against pathogens in the GI tract has been addressed. Generally, probiotics produce numerous chemical antimicrobials to competitively exclude pathogens. Furthermore, probiotics have been considered as potential drug candidates for replacing antibiotics. Pln biosynthesis gene cluster identified in LLY-606 and pc-26 genomes provide us new insights into the exploiting these two strains as probiotics to regulate gut microbiota by resisting pathogens, or to produce effective candidates for antibiotics.

Previous studies demonstrated that high bile salt hydrolase (BSH) activity in the intestine contribute to the declining level of serum cholesterol [9]. At the same time, BSH protect microorganisms from unfavorable bile, which affects their survival in the intestine [52]. Our previous studies have shown that these strains with high BSH activity have well cholesterol-lowering ability *in vitro* [16]. As what was shown in the present study, *bsh* genes existed widely in *L. plantarum* strains. Bile salt hydrolase can not only contribute to a reduction in bile toxicity, but is also associated with the therapeutic effect of lowering serum cholesterol. The presence of BSH genes in all studied *L. plantarum* strains indicated a possible common bile salts or even cholesterol-lowering ability, but these functions was still waited to be confirmed *in vivo*.

By *in vivo* study, the cholesterol-lowering ability was demonstrated in this study. However, whether these probiotic functions were assured by BSH, or by regulating gut microbiota via pln or the carbohydrate fermentation, was not known in this study. But it was no doubt that any of the properties may contribute to its probiotic functions. And the safety, stress response, adhesion ability, antibacterial activity and bile salt hydrolase will add up to their probiotic functions.

In this study, we reported the whole genome of LLY-606 and pc-26. Moreover, intensive genome analysis clarified probiotic properties of LLY-606 and pc-26, including bile salt resistance ability, antibacterial activity, adhesion ability and stress response capabilities. In addition, animal assays in golden hamster demonstrated that administration of LLY-606 or pc-26 reduced total cholesterol level in serum. All these results indicated that they were highly potent probiotic candidates. In this study, a pre-valuation method based on genome sequence for potent probiotic strains were used and probiotic functions were verified in vitro and in vivo.

## 5. Acknowledgments

We would like to thank State Key Laboratory of Agricultural Genomics, Key Lab of Genomics, Ministry of Agriculture and BGI Sanshengyuan Co., Ltd for assistance with implementing experiments. This study was supported by founds from Science, Technology and Innovation Commission of Shenzhen Municipality under grant No. JCYJ20170817151208611 and Shenzhen Dapeng District Government No. KY20180108.

## Supplementary Materials

Figure S1: Pictures of LYY-606 and pc-26 adhere the surface of HT-29 cells; Figure S2: Adhesion efficiency of LYY-606 and pc-26 to HT-29 cells; Figure S3: The location of *bsh* gene in different *L. plantarum* completed genome; Table S1: Clusters of COGs in 56 genomes of *L. plantarum* strains; Table S2: *L. plantarum* strains characterized in this study; Table S3: The prophage regions of *L. plantarum* LYY-606 and *L. plantarum* pc-26; Table S4: Stress-related proteins of *L. plantarum* LYY-606 and *L. plantarum* pc-26; Table S5: The location of *bsh* gene in different *L. plantarum* completed genomes.

## Reference

1. Siezen RJ, Vlieg JETvH. 2011. Genomic diversity and versatility of Lactobacillus plantarum, a natural metabolic engineer. Microbial Cell Factories. 10 (suppl 1): S3.

2. O’Sullivan O, O’Callaghan J, Sangrador-Vegas A, McAuliffe O, Slattery L, Kaleta P, et al. 2009. Comparative genomics of lactic acid bacteria reveals a niche-specific gene set. Bmc Microbiology. 9: 50

3. Mendes-Soares H, Suzuki H, Hickey RJ, Forney LJ. 2014. Comparative Functional Genomics of Lactobacillus spp. Reveals Possible Mechanisms for Specialization of Vaginal Lactobacilli to Their Environment. Journal of Bacteriology. 196: 1458–1470.

4. Martino ME, Bayjanov JR, Caffrey BE, Wels M, Joncour P, Hughes S, et al. 2016. Nomadic lifestyle of Lactobacillus plantarum revealed by comparative genomics of 54 strains isolated from different habitats. Environmental Microbiology. 18: 4974–4989.

5. Molenaar D, Bringel F, Schuren FH, de Vos WM, Siezen RJ, Kleerebezem M. 2005. Exploring Lactobacillus plantarum genome diversity by using microarrays. Journal of Bacteriology. 187: 6119–6127.

6. Dunne C, O’Mahony L, Murphy L, Thornton G, Morrissey D, O’Halloran S, et al. 2001. In vitro selection criteria for probiotic bacteria of human origin: correlation with in vivo findings. American Journal of Clinical Nutrition. 73: 386S–392S.

7. Macklaim JM, Gloor GB, Anukam KC, Cribby S, Reid G. 2011. At the crossroads of vaginal health and disease, the genome sequence of Lactobacillus iners AB-1. Proceedings of the National Academy of Sciences of the United States of America. 108: 4688–4695.

8. Lasek R, Dziewit L, Ciok A, Decewicz P, Romaniuk K, Jedrys Z, et al. 2017. Genome content, metabolic pathways and biotechnological potential of the psychrophilic Arctic bacterium Psychrobacter sp DAB_AL43B, a source and a host of novel Psychrobacter-specific vectors. Journal of Biotechnology. 263: 64–74.

9. Horackova S, Plockova M, Demnerova K. 2018. Importance of microbial defence systems to bile salts and mechanisms of serum cholesterol reduction. Biotechnology Advances. 36: 682–690.

10. An H, Douillard FP, Wang G, Zhai Z, Yang J, Song S, et al. 2014. Integrated Transcriptomic and Proteomic Analysis of the Bile Stress Response in a Centenarian-originated Probiotic Bifidobacterium longum BBMN68. Molecular & Cellular Proteomics. 13: 2558–2572.

11. Hidalgo-Cantabrana C, Sanchez B, Milani C, Ventura M, Margolles A, Ruas-Madiedo P. 2014. Genomic Overview and Biological Functions of Exopolysaccharide Biosynthesis in Bifidobacterium spp. Applied and Environmental Microbiology. 80: 9–18.

12. Corcoran BM, Stanton C, Fitzgerald G, Ross RP. 2008. Life under stress: The probiotic stress response and how it may be manipulated. Current Pharmaceutical Design. 14: 1382–1399.

13. Zankari E, Hasman H, Cosentino S, Vestergaard M, Rasmussen S, Lund O, et al. 2012. Identification of acquired antimicrobial resistance genes. Journal of Antimicrobial Chemotherapy. 67: 2640–2644.

14. Cosentino S, Larsen MV, Aarestrup FM, Lund O. 2013. PathogenFinder - Distinguishing Friend from Foe Using Bacterial Whole Genome Sequence Data. Plos One. 8: e77302

15. Arndt D, Grant JR, Marcu A, Sajed T, Pon A, Liang Y, et al. 2016. PHASTER: a better, faster version of the PHAST phage search tool. Nucleic Acids Research. 44: W16–W21.

16. Sela DA, Chapman J, Adeuya A, Kim JH, Chen F, Whitehead TR, et al. 2008. The genome sequence of Bifidobacterium longum subsp infantis reveals adaptations for milk utilization within the infant microbiome. Proceedings of the National Academy of Sciences of the United States of America. 105: 18964–18969.

17. Schell MA, Karmirantzou M, Snel B, Vilanova D, Berger B, Pessi G, et al. 2002. The genome sequence of Bifidobacterium longum reflects its adaptation to the human gastrointestinal tract. Proceedings of the National Academy of Sciences of the United States of America. 99: 14422–14427.

18. Motherway MOC, Zomer A, Leahy SC, Reunanen J, Bottacini F, Claesson MJ, et al. 2011. Functional genome analysis of Bifidobacterium breve UCC2003 reveals type IVb tight adherence (Tad) pili as an essential and conserved host-colonization factor. Proceedings of the National Academy of Sciences of the United States of America. 108: 11217–11222.

19. Peng C, Li Y, Chen Y, Kong Y, Lin L, Li H, et al. 2017. Screening and comparison of cholesterol-lowing Lactobacillus strains. Zhongguo Weishengtaxixue Zazhi / Chinese Journal of Microecology. 29: 249–255.

20. Koren S, Walenz BP, Berlin K, Miller JR, Bergman NH, Phillippy AM. 2017. Canu: scalable and accurate long-read assembly via adaptive k-mer weighting and repeat separation. Genome Research. 27: 722–736.

21. Chin C-S, Alexander DH, Marks P, Klammer AA, Drake J, Heiner C, et al. 2013. Nonhybrid, finished microbial genome assemblies from long-read SMRT sequencing data. Nature Methods. 10: 563–569.

22. McKenna A, Hanna M, Banks E, Sivachenko A, Cibulskis K, Kernytsky A, et al. 2010. The Genome Analysis Toolkit: A MapReduce framework for analyzing next-generation DNA sequencing data. Genome Research. 20: 1297–1303.

23. Delcher AL, Harmon D, Kasif S, White O, Salzberg SL. 1999. Improved microbial gene identification with GLIMMER. Nucleic Acids Research. 27: 4636–4641.

24. Lowe TM, Eddy SR. 1997. tRNAscan-SE: A program for improved detection of transfer RNA genes in genomic sequence. Nucleic Acids Research. 25: 955–964.

25. Lagesen K, Hallin P, Rodland EA, Staerfeldt H-H, Rognes T, Ussery DW. 2007. RNAmmer: consistent and rapid annotation of ribosomal RNA genes. Nucleic Acids Research. 35: 3100–3108.

26. Burge SW, Daub J, Eberhardt R, Tate J, Barquist L, Nawrocki EP, et al. 2013. Rfam 11.0: 10 years of RNA families. Nucleic Acids Research. 41: D226–D232.

27. Tatusov RL, Natale DA, Garkavtsev IV, Tatusova TA, Shankavaram UT, Rao BS, et al. 2001. The COG database: new developments in phylogenetic classification of proteins from complete genomes. Nucleic Acids Research. 29: 22–28.

28. Overbeek R, Olson R, Pusch GD, Olsen GJ, Davis JJ, Disz T, et al. 2014. The SEED and the Rapid Annotation of microbial genomes using Subsystems Technology (RAST). Nucleic Acids Research. 42: D206–D214.

29. Weber T, Blin K, Duddela S, Krug D, Kim HU, Bruccoleri R, et al. 2015. antiSMASH 3.0-a comprehensive resource for the genome mining of biosynthetic gene clusters. Nucleic Acids Research. 43: W237–W243.

30. Edgar RC. 2004. MUSCLE: multiple sequence alignment with high accuracy and high throughput. Nucleic Acids Research. 32: 1792–1797.

31. Letunic I, Bork P. 2016. Interactive tree of life (iTOL) v3: an online tool for the display and annotation of phylogenetic and other trees. Nucleic Acids Research. 44: W242–W245.

32. Joensen KG, Scheutz F, Lund O, Hasman H, Kaas RS, Nielsen EM, et al. 2014. Real-Time Whole-Genome Sequencing for Routine Typing, Surveillance, and Outbreak Detection of Verotoxigenic Escherichia coli. Journal of Clinical Microbiology. 52: 1501–1510.

33. Grissa I, Vergnaud G, Pourcel C. 2007. CRISPRFinder: a web tool to identify clustered regularly interspaced short palindromic repeats. Nucleic Acids Research. 35: W52–W57.

34. Bertelli C, Laird MR, Williams KP, Lau BY, Hoad G, Winsor GL, et al. 2017. IslandViewer 4: expanded prediction of genomic islands for larger-scale datasets. Nucleic Acids Research. 45: W30–W35.

35. van Heel AJ, de Jong A, Montalban-Lopez M, Kok J, Kuipers OP. 2013. BAGEL3: automated identification of genes encoding bacteriocins and (non-)bactericidal posttranslationally modified peptides. Nucleic Acids Research. 41: W448–W453.

36. Liu C-J, Wang R, Gong F-M, Liu X-F, Zheng H-J, Luo Y-Y, et al. 2015. Complete genome sequences and comparative genome analysis of Lactobacillus plantarum strain 5-2 isolated from fermented soybean. Genomics. 106: 404–411.

37. Ventura M, Canchaya C, Kleerebezem M, de Vos WA, Siezen RJ, Brussow H. 2003. The prophage sequences of Lactobacillus plantarum strain WCFS1. Virology. 316: 245–255.

38. Remus DM, van Kranenburg R, van Swam II, Taverne N, Bongers RS, Wels M, et al. 2012. Impact of 4 Lactobacillus plantarum capsular polysaccharide clusters on surface glycan composition and host cell signaling. Microbial Cell Factories. 11: 149

39. Diep DB, Straume D, Kjos M, Torres C, Nes IF. 2009. An overview of the mosaic bacteriocin pln loci from Lactobacillus plantarum. Peptides. 30: 1562–1574.

40. Tai HF, Foo HL, Rahim RA, Loh TC, Abdullah MP, Yoshinobu K. 2015. Molecular characterisation of new organisation of plnEF and plw loci of bacteriocin genes harbour concomitantly in Lactobacillus plantarum I-UL4. Microbial Cell Factories. 14: 89

41. Alvarez-Sieiro P, Montalban-Lopez M, Mu D, Kuipers OP. 2016. Bacteriocins of lactic acid bacteria: extending the family. Applied Microbiology and Biotechnology. 100: 2939–2951.

42. Lambert JM, Bongers RS, de Vos WM, Kleerebezem M. 2008. Functional analysis of four bile salt hydrolase and penicillin acylase family members in Lactobacillus plantarum WCFS1. Applied and Environmental Microbiology. 74: 4719–4726.

43. Ren J, Sun K, Wu Z, Yao J, Guo B. 2011. All 4 Bile Salt Hydrolase Proteins Are Responsible for the Hydrolysis Activity in Lactobacillus plantarum ST-III. Journal of Food Science. 76: M622–M628.

44. Li P, Li X, Gu Q, Lou X-y, Zhang X-m, Song D-f, et al. 2016. Comparative genomic analysis of Lactobacillus plantarum ZJ316 reveals its genetic adaptation and potential probiotic profiles. Journal of Zhejiang University-Science B. 17: 569–579.

45. Besselink MGH, van Santvoort HC, Buskens E, Boermeester MA, van Goor H, Timmerman HM, et al. 2008. Probiotic prophylaxis in predicted severe acute pancreatitis: a randomised, double-blind, placebo-controlled trial. Lancet. 371: 651–659.

46. Suez J, Zmora N, Zilberman-Schapira G, Mor U, Dori-Bachash M, Bashiardes S, et al. 2018. Post-Antibiotic Gut Mucosal Microbiome Reconstitution Is Impaired by Probiotics and Improved by Autologous FMT. Cell. 174: 1406–1423.

47. Kothari D, Patel S, Kim S-K. 2018. Probiotic supplements might not be universally-effective and safe: A review. Biomedicine & pharmacotherapy = Biomedecine & pharmacotherapie. 111: 537–547.

48. Imperial ICVJ, Ibana JA. 2016. Addressing the Antibiotic Resistance Problem with Probiotics: Reducing the Risk of Its Double-Edged Sword Effect. Frontiers in Microbiology. 7: 1983

49. Jin J, Zhang B, Guo H, Cui J, Jiang L, Song S, et al. 2012. Mechanism Analysis of Acid Tolerance Response of Bifidobacterium longum subsp longum BBMN 68 by Gene Expression Profile Using RNA-Sequencing. Plos One. 7: e50777

50. Moslehi-Jenabian S, Gori K, Jespersen L. 2009. AI-2 signalling is induced by acidic shock in probiotic strains of Lactobacillus spp. International Journal of Food Microbiology. 135: 295–302.

51. Castro-Bravo N, Wells JM, Margolles A, Ruas-Madiedo P. 2018. Interactions of Surface Exopolysaccharides From Bifidobacterium and Lactobacillus Within the Intestinal Environment. Frontiers in Microbiology. 9: 2426

52. Corzo G and Gilliland SE. 1999. Bile salt hydrolase activity of three strains of *Lactobacillus acidophilus*. Journal of Dairy Science. 82: 472–480

